## Supplemental Data for "The inflammasome-activated cytokine IL-1β is targeted for ubiquitylation and proteasomal degradation to limit its inflammatory potential"

**Supplemental Figures**

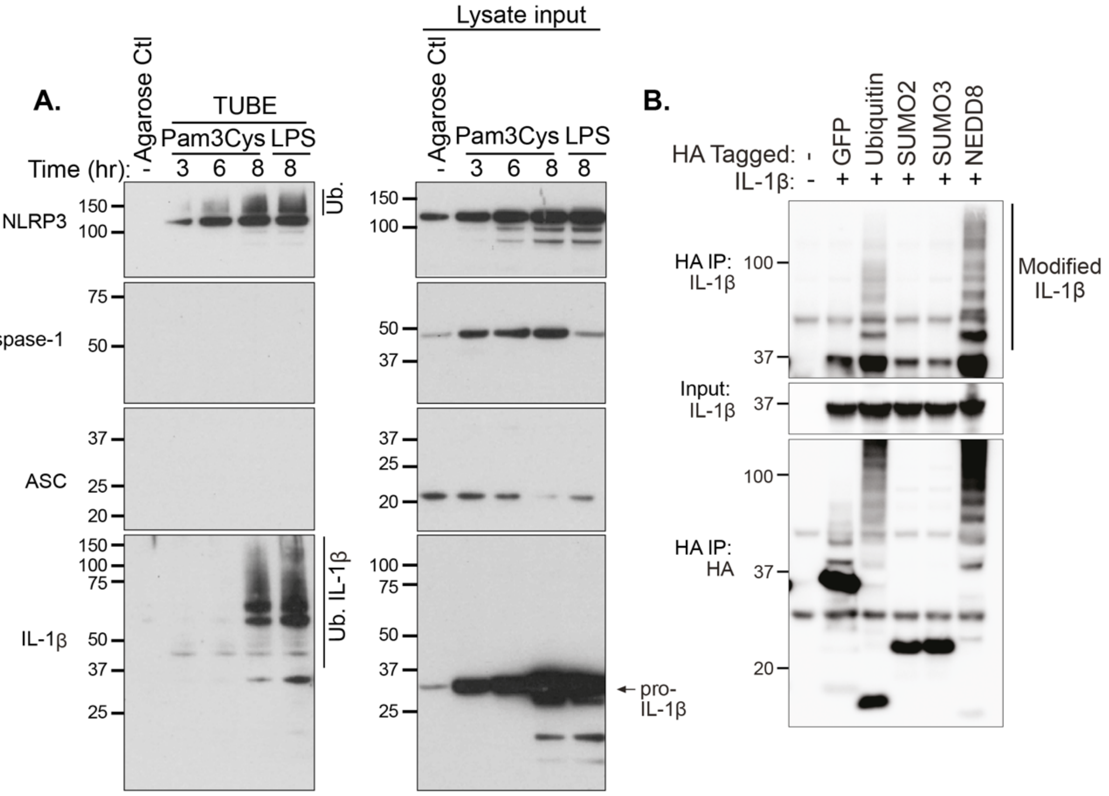


**Supplemental Figure 1. TLR1/2-mediated inflammasome priming triggers IL-1β ubiquitylation**

**A.** BMDMs were primed with Pam3Cys (500 ng/ml) or LPS (50 ng/ml), as indicated, for up to 8 hr and ubiquitylated proteins were isolated from cell lysates using TUBE purification. Immunoblots were performed on cell lysates and purified ubiquitylated proteins (TUBEs) were performed and probed for the specified proteins. Agarose control (ctl) shows the specificity of ubiquitylated protein purification. One of 3 experiments.

**B.** 293T cells were transfected with IL-1β and HA-NEDD8, HA-Ubiquitin, HA-SUMO2, HA-SUMO3 and GFP control, as indicated. HA-tagged proteins were immunoprecipitated and interactions with IL-1β determined by immunoblot.

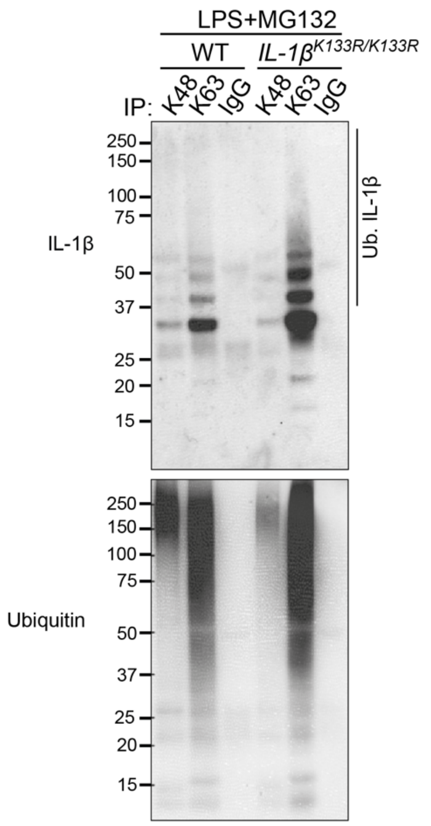


**Supplemental Figure 2. Precursor IL-1β is modified by K48-, K63-linked ubiquitin chains**
WT and *IL-1β* $^{K133R/K133R}$ BMDMs were treated with LPS (50 ng/ml) for 3 hr and then MG132 (20 μM) for a further 2 hr. Ubiquitylated proteins were immunoprecipitated using K48-, K63-specific antibodies (or IgG control) and immunoblotted as indicated. One of 2 experiments.

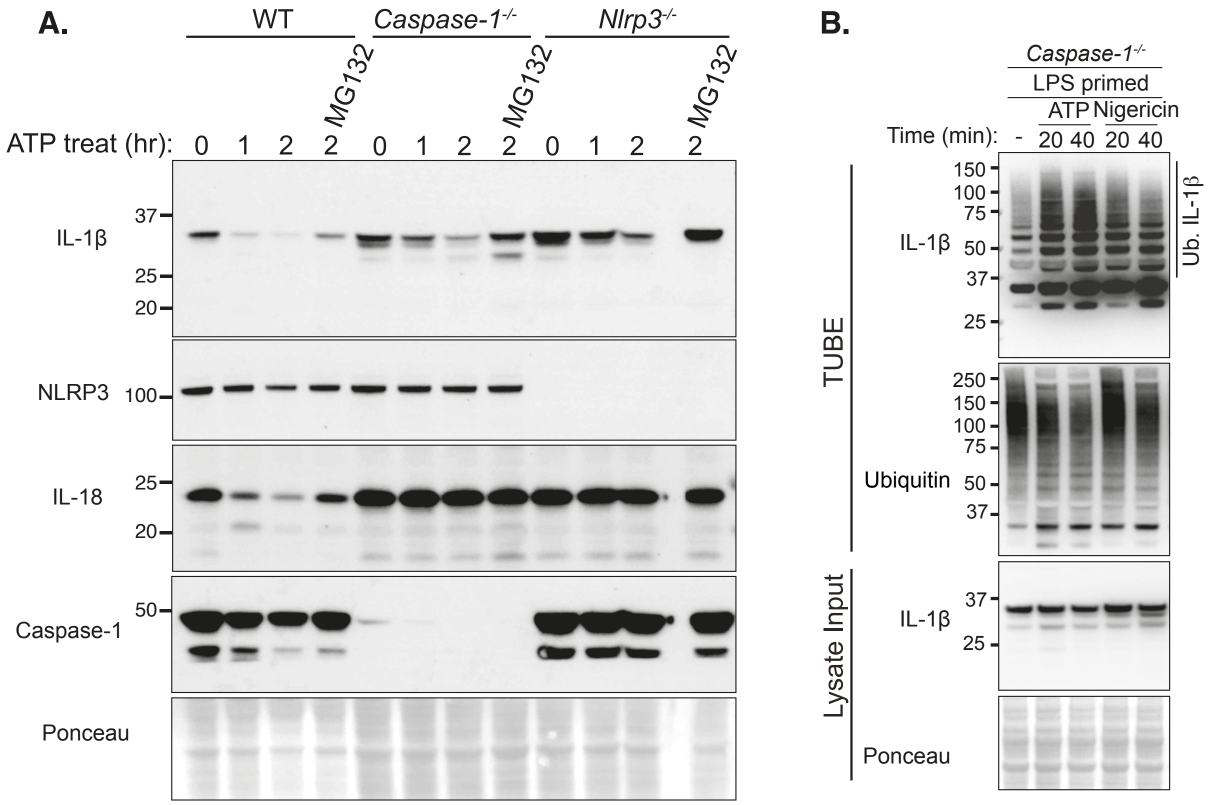


**Supplemental Figure 3. Inflammasome activators can induce proteasomal-mediated IL-1β turnover.**

**A.** WT, *Nlrp3*$^{-/-}$ and *Caspase-1*$^{-/-}$ BMDMs were primed with LPS (50 ng/ml) for 3 hours, and in the last 30 min of priming treated with MG132 (20 μM), as indicated. Cells were treated with ATP (5 mM) for up to 2 hr. Immunoblots were performed on total cell lysates to detect the specified proteins. One of 2 experiments.

**B.** *Caspase-1*$^{-/-}$ BMDMs were generated, LPS primed (100 ng/ml, 3 hr), and then treated with ATP (5 mM) or nigiericin (10 μM) for the indicated times. Ubiquitylated IL-1β was subsequently analyzed by TUBE purification and immunoblotting. 1 of 2 independent experiments.

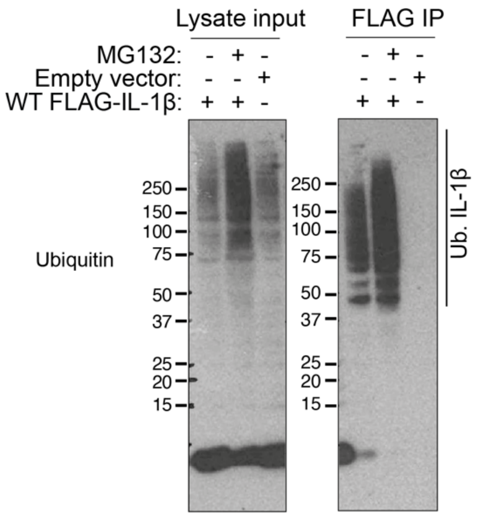


**Supplemental Figure 4. Ubiquitylation of FLAG-IL-1β in 293T cells is enhanced by proteasomal inhibition.**
N-terminal FLAG-IL-1β (pMIGRMCS) or empty vector were transfected into 293T cells and after ~42 hr treated with MG132 (20 μM) for a further 6 hr. FLAG-tagged IL-1β was isolated by FLAG-immunoprecipitation and analyzed by immunoblot, as indicated.

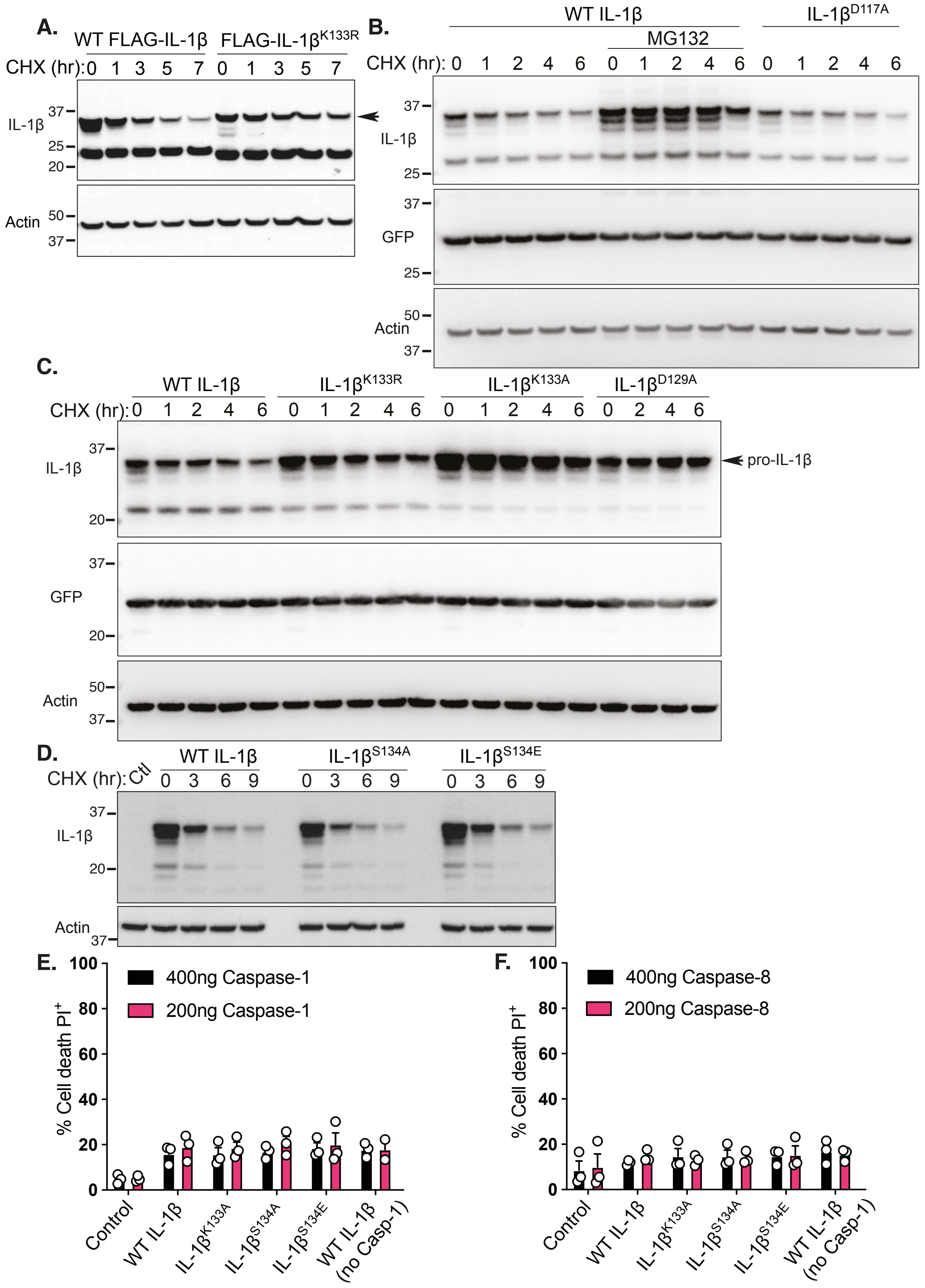


**Supplemental Figure 5. IL-1β K133, but not IL-1β S134, is important for IL-1β stability. A.** FLAG-IL-1β and FLAG-IL-1β$^{K133R}$ cDNA was transfected into 293T cells and after 48 hrs cells were cultured with CHX (20 μg/ml) treatment for up to 7 hr. Immunoblots were performed on total cell lysates for IL-1β protein levels. The arrow indicates pro-IL-1β. Actin is included as a loading control. One of 2 experiments.

**B.** 293T cells expressing WT IL-1β and IL-1β$^{D117A}$ were pre-treated with MG132 (20 μM) for 15 minutes, as indicated, and then treated with CHX (20 μg/ml) for up to 7 hr. Immunoblots were performed on cell lysates for IL-1β and GFP (control for IL-1β plasmid levels). Actin is included as a loading control. One of 2 experiments.

**C.** 293T cells expressing WT IL-1β, IL-1β$^{K133R}$, IL-1β$^{K133A}$ and IL-1β$^{D129A}$ were treated with CHX (20 μg/ml) for up to 6 hr. Immunoblots were performed on cell lysates for IL-1β and GFP (control for IL-1β plasmid levels). Actin is included as a loading control. One of 3 experiments.

**D.** 293T cells expressing WT IL-1β, IL-1β$^{S134A}$ and IL-1β$^{S134E}$ were treated with CHX (20 μg/ml) for up to 9 hr. Immunoblots were performed on cell lysates for IL-1β. Actin is included as a loading control. One of 3 experiments.

**E and F.** 293T cells expressing WT IL-1β, IL-1β$^{K133R}$, IL-1β$^{S134A}$ and IL-1β$^{S134E}$ were co-transfected with 200 and 400 ng of **(D)** caspase-1 or **(E)** caspase-8, as indicated, for 24-48 hr. Cell death was measured by propidium iodide incorporation and flow cytometric analysis. Data are the mean + S.E.M. of 3 independent experiments.

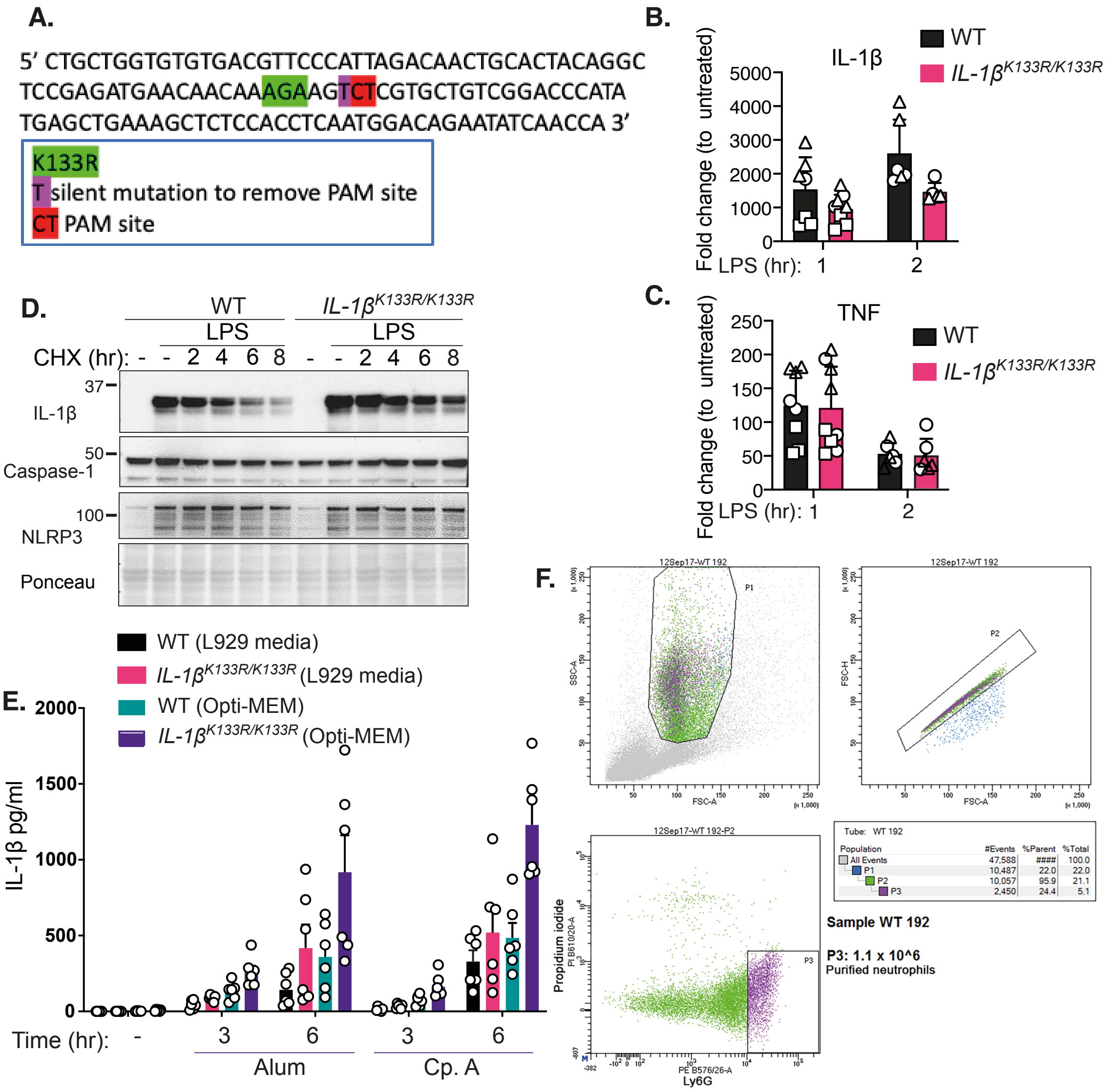


**Supplemental Figure 6. IL-1β K133R protein exhibits increased stability that allows for heightened IL-1β activation.**

**A.** Schema of the CRISPR mouse sgRNA targeting oligonucleotide used for generating the IL-1β K133R knock-in mouse.

**B and C.** WT and *IL-1β* $^{K133R/K133R}$ BMDMs were treated with or without LPS (50 ng/ml) for up to 2 hr. **(B)** IL-1β and **(C)** TNF mRNA levels were analyzed by RT-qPCR. Data are the mean + SD. Individual mice/replicates are shown by symbols from 3 pooled independent experiments.

**D.** WT and *IL-1β* $^{K133R/K133R}$ BMDMs were treated with **(D)** LPS (50 ng/ml) for 3 hr and then treated with CHX (20 µg/ml) for up to 8 hr. Cell lysates were analyzed for the indicated proteins by immunoblot. One of 4 experiments.

**E.** WT and *IL-1β* $^{K133R/K133R}$ BMDMs were cultured in either L929-conditioned complete DMEM or Opti-MEM media, primed with LPS (50 ng/ml) for 3 hr and treated with Alum (300 µg/ml) or Smac-mimetic (Compound A, 500 nM) for up to 6 hr. Cell supernatants were analyzed for levels of IL-1β by ELISA. Data are mean + SEM, n = 3 mice, representative of 1 of 3 experiments.

**E.** Gating strategy for bone marrow neutrophil purification.

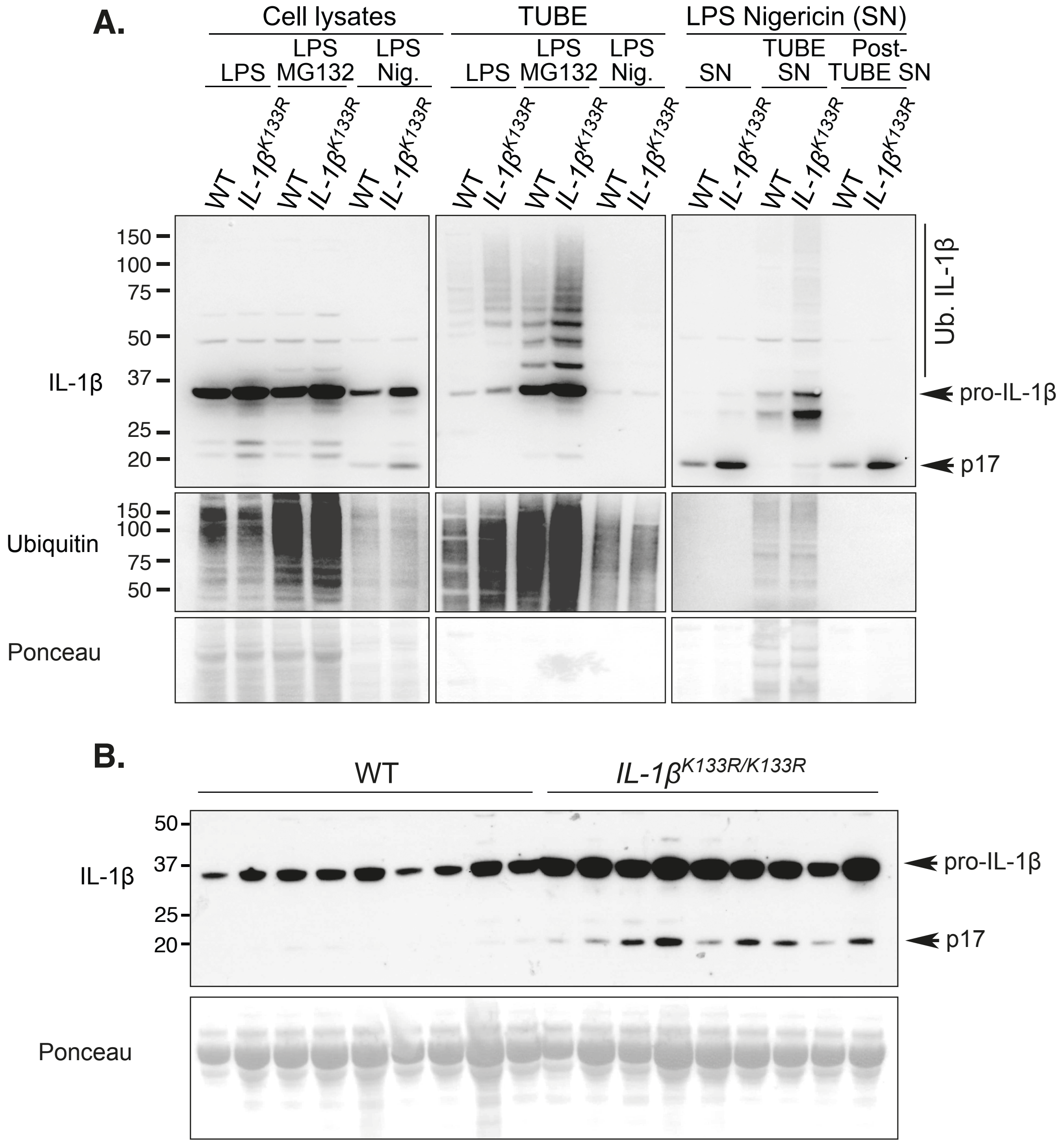


**Supplemental Figure 7. LPS-induced IL-1β is increased in *IL-1β*$^{K133R/K133R}$ mice *in vivo*.**

**A.** WT and *IL-1β*$^{K133R/K133R}$ BMDMs (10 million cells) were primed with LPS (50 ng/ml) for 3 hr and then treated with MG132 (20 μM) for a further 2 hr, or treated with nigericin (20 μM) in Opti-MEM media for 40 min. Ubiquitylated proteins were purified from cell lysates and 5 ml of supernatants using TUBEs. As specified, cell lysates, supernatants and TUBE enriched proteins (including TUBE-depleted supernatants) were analyzed by immunoblot. Note, equivalent amounts of supernatant and post-TUBE supernatant were analyzed to allow a direct comparison. One of 2 experiments.

**B.** WT and *IL-1β*$^{K133R/K133R}$ mice were injected intra-peritoneally with 100 μg of LPS. After 2 hr, peritoneal fluid was harvested and analyzed by immunoblot. Each lane represents an individual mouse. n= 9 mice per genotype.
